# Brain Abnormalities Associated with Self-Injurious Thoughts and Behaviors: A Meta-Analysis of Neuroimaging Studies

**DOI:** 10.1101/526525

**Authors:** Xieyining Huang, Kelly Rootes-Murdy, Diana M. Bastidas, Derek E. Nee, Joseph C. Franklin

## Abstract

Self-injurious thoughts and behaviors (SITBs) have long been believed to result in part from brain abnormalities. This meta-analysis aims to evaluate whether the extant literature justifies any definitive conclusions about whether and how SITBs may be associated with aberrant findings. Sixty studies published through June 1st, 2017 that compared individuals with and without SITBs were included, resulting in 734 coordinates. A pooled meta-analysis assessing for general risk for SITBs indicated a lack of convergence on structural abnormalities. Functional abnormalities in the precuneus/posterior cingulate cortex (PCC), temporal-parietal junction, and rostral-lateral prefrontal cortex were significant using multi-level kernel density analysis but nonsignificant using activation-likelihood estimation. Separate analyses for types of SITBs suggested that deliberate self-harm might be associated with abnormalities in the precuneus/PCC. Some moderator effects were detected. Overall, the meta-analytic evidence was not robust. More studies are needed to reach definitive conclusions about whether SITBs are associated with brain abnormalities.

---

Self-injurious thoughts and behaviors (SITBs) are major public health concerns (Nock et al. 2008; Nock 2010; Centers for Disease Control and Prevention, 2016). Researchers have proposed that SITBs result in part from biological causes in brain systems, and many studies have linked SITBs to specific brain abnormalities (Lutz, Mechawar, & Turecki, 2017). Multiple researchers have proposed that examining the “suicidal brain” will improve our understanding of SITBs, accelerate the discovery of useful biomarkers, improve the identification of high-risk individuals, and inform therapeutic and pharmacological targets for treatment (Desmyter, van Heeringen, & Audenaert, 2011; Jollant, Lawrence, Olié, Guillaume, & Courtet, 2011; Mann, 2003; Turecki, 2014). However, the results of neuroimaging studies examining SITBs vary considerably, making it difficult to synthesize evidence on specific brain abnormalities based on qualitative reviews. As such, this study aims to quantitatively summarize existing brain imaging research on SITBs.

Among individuals who experience SITBs, recent neuroimaging studies have found abnormalities in multiple brain regions associated with psychological traits that confer risk for SITBs. For instance, structural alterations have been reported in the ventrolateral prefrontal cortex, dorsal prefrontal cortex, and orbitofrontal cortex among suicide attempters, and these abnormalities were considered to reflect deficient cognitive control and impaired decision-making in the context of intense emotions (Ding et al., 2015). Other studies reported similar findings and conclusions with regard to the amygdala (Monkul et al., 2007), the anterior cingulate cortex (Wagner et al., 2011), and the midbrain/pons (Osuch, Ford, Wrath, Bartha, & Neufeld, 2014), and their associations with traits such as impulsivity, heightened stress reactivity, emotion dysregulation, and pain perception. Given these findings, researchers have proposed that future studies should focus on examining imaging endophenotypes associated with vulnerability to SITBs (Courtet, Gottesman, Jollant, & Gould, 2011). Some researchers further encouraged efforts to identify a biosignature of suicide and develop a biologically-based model of suicide risk (Oquendo et al., 2014; Sudol & Mann, 2017).

Despite evidence linking brain abnormalities with vulnerability to SITBs, neuroimaging studies often yield different results. Most brain imaging studies that examine differences between SITB and non-SITB populations detect some kind of abnormality (though some studies have reported null effects [Ballard et al. 2015; Richard-Devantoy et al. 2016]), but the same abnormality is rarely detected across multiple studies. For example, multiple studies have found decreased gray matter volumes in suicide attempters, but the location of these abnormalities has varied. One study found this alteration in the left superior temporal lobe and left orbitofrontal cortex (Aguilar et al., 2008), another study solely discovered this difference in the insula and posterior cingulate regions (Hwang et al., 2010), and a third study found broader altered regions including the dorsolateral prefrontal cortex, orbitofrontal cortex, and the parieto-occipital cortex (Benedetti et al., 2011). Given these inconsistent findings, does the existing literature justify any definitive conclusions about abnormal brain structure or function among people with a history of SITBs?

According to multiple qualitative reviews of portions of this literature, the literature does justify such conclusions. Notably, however, different reviews reach different conclusions. For instance, reviews have concluded, based on heterogeneous mixes of approximately 10-20 studies each, that SITBs are associated with abnormalities in the ventrolateral, orbital, dorsomedial, dorsolateral, and ventromedial prefrontal cortices, the anterior cingulate gyrus, the amygdala, white matter connection, and gray matter volume (e.g., Heeringen, Bijttebier, & Godfrin, 2011; Jollant et al., 2011; Mann, 2003; Martin, Zimmer, & Pan, 2015; Zhang, Chen, Jia, & Gong, 2014). These reviews discussed studies examining a wide range of SITBs (e.g., suicide ideation, intent, lethality, NSSI) and included diverse imaging methods (e.g., CT, MRI, SPECT, PET, fMRI, DTI) and neuropsychological tasks (e.g., viewing emotional faces, Iowa Gambling Task, verbal fluency task, Stroop task). The samples included in these reviews were also diverse, with psychiatric diagnoses of Schizophrenia, Borderline Personality Disorder, Bipolar Disorder, Major Depressive Disorder, and various affective disorders. The inconsistent conclusions across these reviews leaves open many questions about potential brain structure and functional abnormalities among people with a history of SITBs.

A quantitative meta-analytic review may help to resolve some of these questions. With qualitative methods, it is difficult to accurately weight findings and there is often a tendency to overemphasize positive findings (Borenstein, Hedges, Higgins, & Rothstein, 2009). Quantitative meta-analysis has the advantages of increasing precision in estimating effects, boosting power by combining samples, and evaluating the effects of moderators (Walker, Hernandez, & Kattan, 2008). Compared to narrative reviews, however, fewer meta-analyses exist on this topic. Among the efforts, van Heeringen and colleagues (2014) synthesized 6 structural and 6 functional imaging studies and concluded that suicide attempters have reduced volumes of the rectal gyrus, superior temporal gyrus and caudate nucleus, and increased reactivity of the anterior and posterior cingulate cortices. However, 6 studies per analysis is likely to be too few to draw meaningful conclusions (Eickhoff et al., 2016). Hence, a more comprehensive treatment is needed.

The primary goal of the present study was to conduct a more comprehensive meta-analytic review of the SITB brain imaging literature. Accordingly, the meta-analysis included 60 studies, an improvement over previous reviews. This analysis will help to determine whether the extant literature justifies any definitive conclusions about brain structure or function abnormalities among people with a history of SITBs. This is important for three reasons. First, as reviewed above, there are many inconsistent findings in the literature. Second, the existing meta-analyses were largely underpowered. Guidelines suggest that at least 20 experiments are needed to reliably detect moderate effect sizes in Activation Likelihood Estimation, a common method for performing coordinate-based neuroimaging meta-analyses (Eickhoff et al., 2016). Without sufficient power, true positive findings might be observed in different brain regions even for identically performed studies (Poldrack et al., 2017). Therefore, a meta-analysis with a broader scope and larger power is needed to overcome the limitations and reveal a reliable pattern in the literature. Third, recent comprehensive meta-analyses in other domains have surprisingly found that the existing literature does not justify definitive conclusions about brain imaging differences. For example, across 57 studies, a meta-analysis (Müller et al., 2017) found no evidence for brain imaging abnormalities among depressed individuals. These findings did not prove that there are no neural correlates of depression; rather, they showed that the extant literature did not yet justify any definitive conclusions about the neural correlates of depression and clarified directions for future research aimed at investigating these correlates. Given the inconsistency of findings from the SITB brain imaging literature, the present meta-analyses may obtain similar findings and serve a similar function.

The present study represents one of the largest efforts to synthesize the evidence on structural and functional brain abnormalities associated with SITBs. We had three major aims. First, a pooled analysis on studies examining the neural correlates of any type of SITBs was conducted to test whether certain brain changes were associated with general vulnerabilities to SITBs. Second, separate meta-analyses were conducted for each type of SITBs to examine whether a unique brain aberration exists for different types of SITBs. Third, analyses were conducted to test whether differences in study designs might moderate the findings. The results of this meta-analysis will help to summarize knowledge about the association between SITBs and brain structures and functions, and may serve as a foundation for future work in this area.

## Methods and Materials

### Literature Search and Inclusion Criteria

We identified relevant articles using a range of search terms through June 1st, 2017 using PubMed, PsycInfo, and Google Scholar. We intentionally chose a large number of search terms to increase likelihood of identifying relevant articles that may have been missed otherwise. Search terms included variants of “suicide,” “self-injury,” “self-harm,” “self-directed violence,” “self-mutilation,” “deliberate self-harm (DSH),” and “nonsuicidal self-injury (NSSI)” crossed with variants of “computerized tomography (CT),” “magnetic resonance imaging (MRI),” positron emission tomography (PET),” “single-photon emission computed tomography (SPECT),” “diffusion tensor imaging (DTI),” “magnetic resonance spectroscopy (MRS),” “neuroimaging,” “gray matter,” “white matter,” “serotonin,” “dopamine,” “gamma-aminoburtyric acid,” noradrenaline,” “norepinephrine,” and “brain.”

Inclusion required that studies (1) include at least one group of which all individuals exhibit SITBs; (2) include at least one control group; (3) conduct whole-brain analyses; and (4) provide standardized coordinates. The first inclusion criterion is to ensure that the findings are uniquely associated with SITBs instead of general psychopathology, with the second criterion ensuring that each study provided a benchmark for comparison. The third criterion is to prevent Region of Interest (ROI) analyses from biasing the meta-analytic results. Although ROI analyses provide valuable information about the neural correlates of SITBs, they violate the assumption that each voxel has an equal chance of being activated, thus biasing the meta-analytic results toward convergence on the ROIs (Müller et al., 2018). This inclusion criterion is consistent with other meta-analyses of brain imaging studies (Müller et al., 2017; Norman et al., 2016).

A total of 803 unique papers were identified through database searching. A total of 60 studies were retained in the present study, yielding a total sample size of 3,132 unique participants and 734 unique coordinates (see Figure S1 for PRISMA flowchart, Supplement 1 for a list of included studies, and Table S1 for description of the studies and contrasts).

### Data Extraction

We extracted the following information from each study: (1) sample size; (2) imaging techniques; (3) type of SITBs; (4) type of control groups (i.e., self-injurious, psychiatric, healthy controls); (5) psychiatric diagnoses; (6) Montreal Neurological Institute (MNI) or Talaraich coordinates; (7) study paradigm (i.e., resting-state versus task-based, with tasks further divided into cognitive versus affective tasks); (8) sample age, and (9) sample medication status.

#### Sample size

We extracted the sample size associated with each contrast.

#### Imaging Techniques

Consistent with previous reviews and meta-analysis, we included studies with a range of imaging techniques (e.g., Jollant et al., 2011; Mann, 2003; van Heeringen, Bijttebier, & Godfrin, 2011; Martin, Zimmer, & Pan, 2015; Zhang et al., 2014; van Heeringen et al., 2014). The types of imaging techniques were determined from each study: Magnetic Resonance Imaging (MRI), functional Magnetic Resonance Imaging (fMRI), Diffusion Tensor Imaging (DTI), Positron Emission Tomography (PET), Single-Photon Emission Computed Topography (SPECT), and Magnetic Resonance Spectroscopy (MRS).

#### Type of SITBs

We adhered to the terminology proposed by Nock (2010) and categorized SITBs examined by each study into: suicide ideation, suicide plan, suicide attempt, suicide death, and nonsuicidal self-injury (NSSI). When a study examined deliberate self-injuries of which the intent to die was unclear, we labeled the type of SITBs as self-harm. When a study examined suicide attempt and other suicidal behaviors (e.g., interrupted attempt, aborted attempt) together, the study was considered to have examined all suicidal behaviors. We intentionally included all types of SITBs to conduct a pooled meta-analysis as well as finer-grained analyses to test whether certain brain abnormalities are associated with general risk for SITBs or only specific types of SITBs.

#### Type of Control Groups

A control group was considered a self-injurious control if participants were selected based on prior or current SITBs (e.g., suicide ideation, NSSI). A control group was coded as a psychiatric control if participants were drawn because they met certain clinical conditions (e.g., a psychiatric diagnosis, a score above the clinical threshold on a measure). When neither eligibility criteria were set by the study, the control group was considered as a healthy control. This code was intended to test whether the stringency of control group might have contributed to the diverse findings in the literature.

#### Psychiatric diagnoses

Given that some evidence suggests that the brain abnormalities associated with SITBs might vary depending on the psychiatric diagnoses (Giakoumatos et al., 2013; Sudol & Mann, 2017), we coded for the primary psychiatric diagnoses of the samples.

#### Montreal Neurological Institute (MNI) or Talairach Coordinates

Whenever provided, MNI or Talairach coordinates were directly extracted for each contrasts from the studies. If a study did not specify whether they provided MNI or Talairach coordinates, the type of coordinates were inferred based on the statistical software used by the authors.

#### Study Paradigm

Following convention of prior reviews and meta-analyses (e.g., van Heeringen et al., 2011; Jollant et al., 2011; Mann, 2003; Martin et al., 2015; van Heeringen et al., 2014; Zhang et al., 2014), we included studies using a wide range of study paradigms. To estimate and control for differences between studies, we first categorized each contrast based on whether they were obtained via resting-state or task-based paradigms. We then categorized task-based paradigms into cognitive versus affective tasks. Even though it was our original intention to code for specific tasks (e.g., Stroop task, Iowa Gambling Tasks) and to test whether they moderate the findings, we were unable conduct such analyses due to heterogeneity in the literature. Even though non-neuroimaging meta-analyses have analyzed these tasks when there were at least three studies using the same task (e.g., Richard-Devantoy et al., 2014), guidelines suggest a minimum of 20 experiments in each category for coordinate-based meta-analyses (Eickhoff et al., 2016). As such, we were unable to produce finer-grained categorizations.

#### Sample Age

The mean sample age was extracted from each contrast. We also categorized sample age into adult, adolescent, and mixed samples. A sample was coded as adult if all the participants were at least 18 years old. It was coded as adolescent if all the participants were under the age of 18. When a sample included both adult and adolescent participants, it was coded as mixed.

#### Medication Status

To assess for potential moderator effects, samples were coded into the following categories based on participants’ psychiatric medication status: none medicated, some medicated, or all medicated.

### Statistical Analysis

The goal of this meta-analysis was to identify brain areas that were consistently related to SITBs. We addressed this goal using two methods of coordinate-based meta-analysis (CBMA): Activation Likelihood Estimation (ALE; Turkeltaub et al. 2002, 2012; Eickhoff et al. 2009) and Multi-level Kernel Density Analysis (MKDA; Wager et al. 2007). Ideally, meta-analysis of neuroimaging data would be performed on statistical maps. Unfortunately, such maps are rarely available despite current efforts to create map repositories (Gorgolewski et al., 2015, 2016). In lieu of such maps, CBMA infers statistical maps based upon locations of statistical local maxima (i.e. peaks). Then, spatial-consistency among the inferred statistical maps is assessed. As detailed below, the two methods employed here differ in how the inferred statistical maps are calculated. The use of two different meta-analytic procedures was to ensure that the results did not depend on methodological specifics. In both cases, statistical significance was determined using cluster-level family-wise error (FWE) correction to control for multiple comparisons, as has been recommended for CBMA (Eickhoff et al., 2016). Cluster extents were determined at a height threshold of *p* < 0.001 using Monte-Carlo permutation methods (Eickhoff, Bzdok, Laird, Kurth, & Fox, 2012; Wager et al., 2007).

Meta-analyses followed a tree approach to assess structural (Figure 1) and functional (Figure 2) abnormalities between populations with and without SITBs. First, to assess for general brain abnormalities that might predispose all individuals to all types of SITBs, we conducted pooled analyses that included all SITBs and all types of sample populations. Second, to test for specific brain changes associated with specific types of SITBs, we conducted separate analyses for each type of SITBs when power allowed. Lastly, to test for potential moderator effects, we conducted more granular analyses based on control group type, psychiatric diagnoses, task type, and medication status, provided sufficient power. We subscribed to the power guidelines proposed by Eickhoff and colleagues (2016), which required a minimum of 20 experiments in each category for ALE. Therefore, in our main report, we focused on those analyses that had 20 or more experiments. For completeness, we reported analyses with at least 10 experiments in Table S2, which had been an earlier criterion (Eickhoff & Bzdok, 2013).

**Figure 1.**
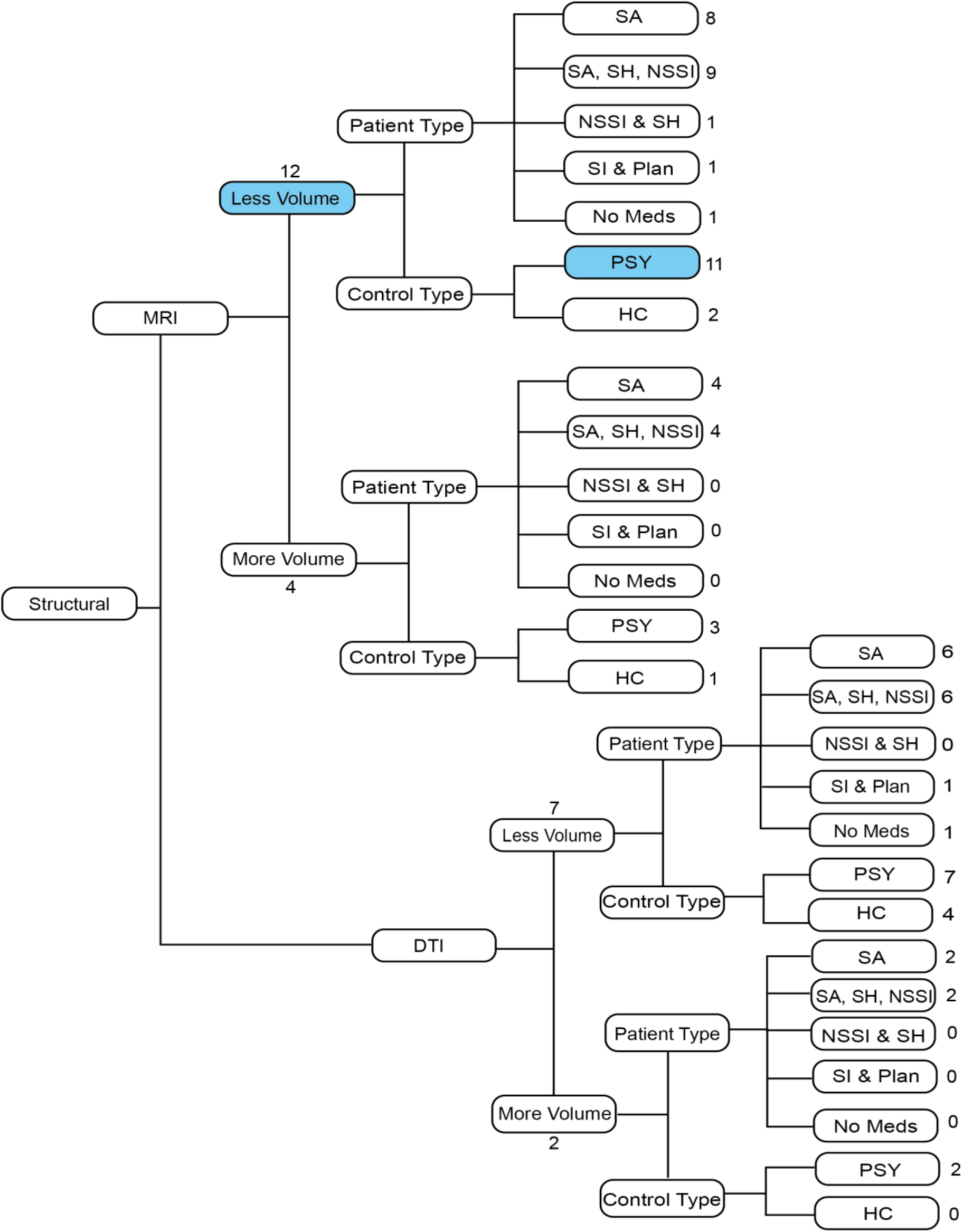
Breakdown of Structural Studies.

*Note*. Contrasts with 10 or more experiments are highlighted in blue. There were no structural contrasts with 20 or more experiments. There were also no significant findings from the structural contrasts in either the MKDA or ALE analyses. HC: healthy controls; PSY: matched psychiatric populations; SITB: all self-injurious thoughts and behaviors; SA: suicide attempts; SH: self-harm; NSSI: non-suicidal self-injury; SI: suicidal ideation; DTI: diffusion tension imaging; MRI: magnetic resonance imaging.

**Figure 2.**
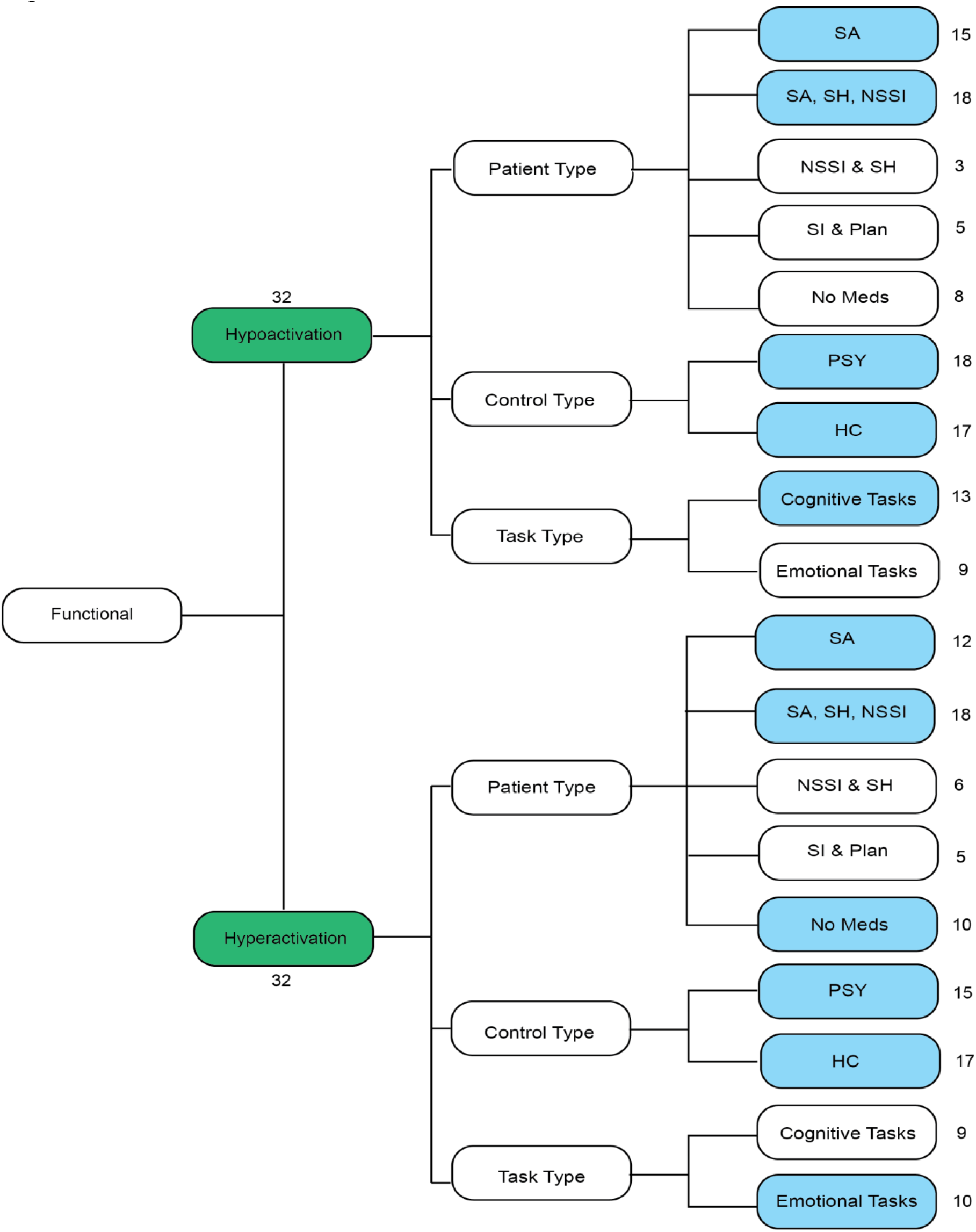
Breakdown of Functional Studies.

*Note*. All possible contrasts from the meta-analysis are listed above. Those contrasts that contained 20 or more experiments are highlighted in green. Those contrasts that contained 10 or more experiments are highlighted in blue. Number of contrasts are listed next to each contrast box. HC: healthy controls; PSY: matched psychiatric populations; SITB: all self-injurious thoughts and behaviors; SA: patients with suicide attempts; SH: self-harm; NSSI: non-suicidal self-injury; SI: suicidal ideation; DTI: diffusion tension imaging; MRI: magnetic resonance imaging.

#### Activation Likelihood Estimation (ALE): Overview

ALE was performed using GingerALE software version 2.3 (Eickhoff et al., 2009; Turkeltaub et al., 2012, 2002). For each experiment, ALE computes “modeled activation” maps that indicate the probability that a given voxel was “activated.” (Although “activated” suggests a functional change, the same logic can be applied to structural data. We use the term “activated” for convenience and consistency with the methodological reports). ALE assumes that each peak represents a broader activation cluster and that the exact location of each peak/cluster is uncertain. Therefore, each peak is convolved with a Gaussian kernel to form a Gaussian probability density. The kernel has a fixed area-under-the-curve, but the full-width/half-maximum (FWHM) varies according to the sample size of the experiment, with the FWHM values empirically determined (Eickhoff et al., 2009). This results in narrower, higher amplitude peak densities for large sample sizes reflecting greater certainty, and broader, lower amplitude peak densities for small sample sizes reflecting less certainty. To control for the fact that studies vary in whether or not sub-peaks within a cluster are reported, we used the non-additive approach that assigns the maximal density amplitude to a voxel that is activated across multiple clusters (Turkeltaub et al., 2012). ALE values are computed for each voxel via the voxel-wise union of the modeled activation maps. Observed ALE values are then compared to a randomly permuted null distribution to determine significance (Eickhoff et al., 2012). Cluster extents were determined at a height threshold of *p* < 0.001 using previously recommended Monte-Carlo permutation methods of 1000 permutations (Eickhoff et al., 2012).

#### Multi-level Kernel Density Analysis (MKDA): Overview

MKDA (Wager et al., 2007) differs from ALE in terms of the kernel that is convolved with each activation peak. MKDA uses a spherical kernel whose radius is determined by the analyst, whereas ALE uses a Gaussian kernel whose FWHM is empirically determined. At first blush, an empirically determined kernel extent may seem superior to an arbitrarily assigned kernel extent. However, the empirically determined FWHM is based upon data from 21 healthy participants performing a single task with BOLD imaging (Eickhoff et al., 2009). Whether the extents observed there generalize to different populations, imaging modalities, and tasks is unclear. Therefore, the ability to freely choose a kernel extent in MKDA offers assurance that significant/non-significant results are not due to this limitation.

We conducted analyses with kernel radii at 10mm and 15mm, which has been previously recommended (Wager, Jonides, & Reading, 2004; Wager et al., 2007). For each study, each peak was convolved with the kernel to create a comparison indicator map. The map has values of either 1 (‘a study activated near this voxel’) or 0 (‘a study did not activate near this voxel’). Similar to the non-additive approach to ALE, the nesting of peaks within studies allows that no one study can disproportionally contribute to the significant findings. Each map is weighted by the product of the square root of the study sample size. The weighted average of these maps is then compared to a randomly permuted null distribution to determine significance. Cluster extents were determined at a height threshold of *p* < 0.001 using the previously recommended Monte-Carlo permutation methods of 5000 permutations (Wager et al., 2007).

#### Sample Size Determination

Both ALE and MKDA weight studies by sample size. Both methods were developed with one-sample tests in mind and thus the weighting procedures assume a one-sample *n*. Here, we are explicitly focused on two-sample tests. To provide an equivalent one-sample *n* we used the equation (*n*_*1*_ × *n*_*2*_)/ (*n*_*1*_ + *n*_*2*_) following prior guidelines (Tench, Tanasescu, Constantinescu, Auer, & Cottam, 2017).

#### Spatial Distributions

Both ALE and MKDA use Monte-Carlo procedures to determine the null distribution. By default, both ALE and MKDA restrict random permutation to gray matter. However, this procedure is not appropriate for analyses that are expected to produce results in white matter (e.g. DTI). Therefore, for MKDA analyses of DTI data, we used a white matter, rather than gray matter mask.

## Results

### Descriptive Statistics

Most of the contrasts were yielded from studies using fMRI (59.13%), followed by structural MRI (18.53%), SPECT (13.22%), DTI (6.13%), and PET (3.00%). Regarding types of SITBs, 44.28% of the contrasts examined suicide attempt, with the rest studying self-harm regardless of suicidal intent (17.03%), NSSI (12.94%), suicide death (11.04%), suicide ideation and plan (9.67%), all suicidal behaviors (2.32%), and suicide risk (2.04%). More than half of the contrasts (61.51%) adopted task-based paradigms (affective tasks: 54.70%; cognitive tasks: 45.30%).

Among the 60 studies included in this meta-analysis, the median sample size was 45 (mean = 58.15, SD = 32.16). Majority of the samples (70.00%) were adult participants, followed by mixed adult and adolescent (15.00%), and child or adolescent (13.33%). One study only included elderly participants (1.67%). The mean sample age was 31.05 years old (SD = 12.37). Half of the samples (50.00%) included some participants with psychiatric medication at the time of the study, with 21.67% of the samples only including unmedicated participants, and 10.00% of the samples only including medicated participants. The rest of the studies (18.33%) did not mention participants’ medication status. Regarding control group type, about half of the contrasts (52.72%) used psychiatric controls, with the rest using healthy controls (43.05%) and self-injurious controls (4.22%; e.g., suicide attempters compared with suicide ideators). Regarding psychiatric diagnoses, about half of the studies (45.00%) required participants to meet criteria for Major Depressive Disorder (MDD), followed by no specific psychiatric diagnoses required (11.67%), Borderline Personality Disorder (8.33%), Schizophrenia (8.33%), Bipolar Disorder (6.67%), unspecified depressive disorder (6.67%), Bipolar Disorder or MDD with psychotic features (3.33%), unipolar or bipolar depression (3.33%), MDD or Posttraumatic Stress Disorder (PTSD) or Traumatic Brain Injury (1.67%), various mood disorders (1.67%), PTSD (1.67%), and Schizophrenia or Schizophreniform Disorder (1.67%).

### Meta-Analytic Results

#### Overall Meta-Analyses

To examine the general brain abnormalities associated with SITBs, the meta-analyses first considered structural and functional data across all types of SITBs and populations.

##### Structural Imaging Studies

No structural analysis featured 20 or more experiments. Although more than 10 experiments reported reduced gray matter volumes in SITBs, neither ALE nor MKDA observed a consistently significant result.

##### Functional Imaging Studies

No significant results were observed with any ALE analyses at either the standard 20 experiment criterion or the relaxed 10 experiment criterion. However, MKDA with a 10mm kernel requiring 20 experiments did reveal hyperactivation in SITBs at the precuneus/posterior cingulate cortex (PCC; Figure 3 and Table 1), which was also observed when the kernel was increased to 15mm. The temporal-parietal junction (TPJ) was also hyperactivated in SITBs at 15mm (Figure 3A). In addition, hypoactivation in SITBs was observed in the rostral-lateral prefrontal cortex (RLPFC) at 15mm (Figure 3B and Table 1). A corresponding RLPFC effect was observed at 10mm, but only when thresholding at the voxel, rather than the cluster-level.

**Table 1:**
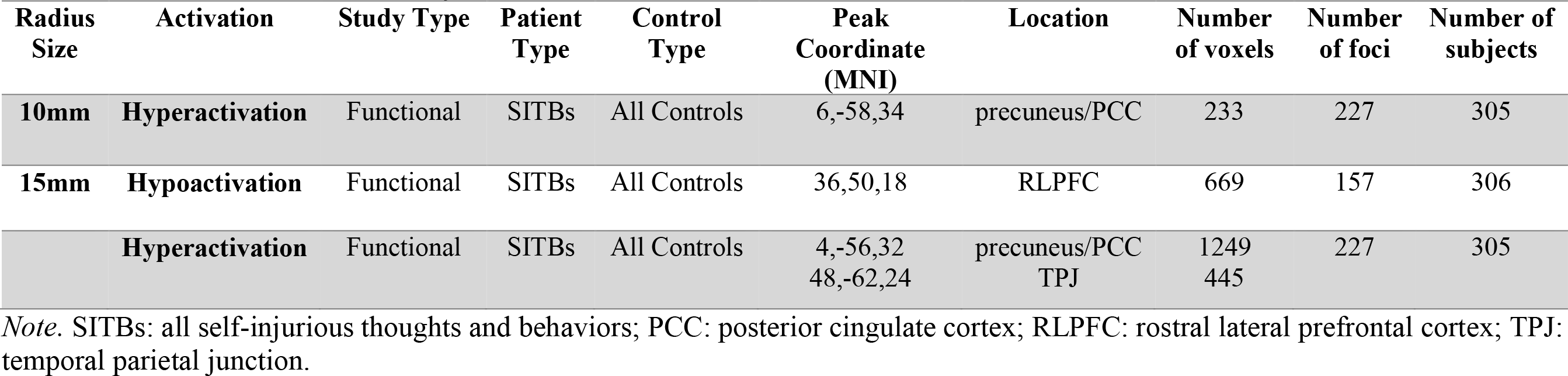
MKDA Coordinate Sites by Contrast for 10mm and 15mm Radii.

| Radius Size | Activation | Study Type | Patient Type | Control Type | Peak Coordinate (MNI) | Location | Number of voxels | Number of foci | Number of subjects |
| --- | --- | --- | --- | --- | --- | --- | --- | --- | --- |
| **10mm** | **Hyperactivation** | Functional | SITBs | All Controls | 6,-58,34 | precuneus/PCC | 233 | 227 | 305 |
| **15mm** | **Hypoactivation** | Functional | SITBs | All Controls | 36,50,18 | RLPFC | 669 | 157 | 306 |
|  | **Hyperactivation** | Functional | SITBs | All Controls | 4,-56,32<br>48,-62,24 | precuneus/PCC<br>TPJ | 1249<br>445 | 227 | 305 |

*Note.* SITBs: all self-injurious thoughts and behaviors; PCC: posterior cingulate cortex; RLPFC: rostral lateral prefrontal cortex; TPJ: temporal parietal junction.

**Figure 3.**
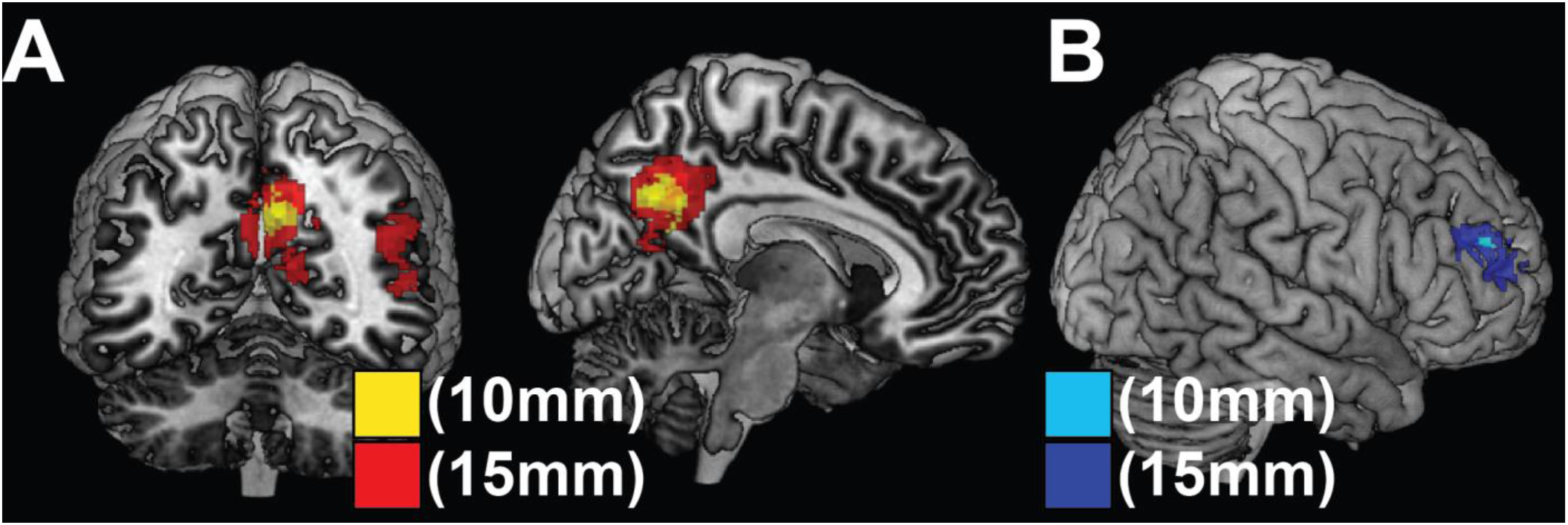
Results of Functional Imaging Studies from MKDA Method at 10mm and 15mm radii. **A)** Significant results for SITB > Controls using MKDA at 10mm (yellow) and at 15mm (red). **B)** Significant results for SITB < Controls using MKDA at 10mm (light blue) and 15mm (dark blue). Results at 10mm survived voxel-level, but not cluster-level correction for multiple comparisons.

##### Functional Inference

To infer the functional consequences of observed significant results, we performed functional specialization classification. Following the procedures of de la Vega and colleagues (2017), the psychological concepts that activate each significant cluster at radius 15mm were inferred via machine learning classification on the NeuroSynth database. The classification determines the extent to which psychological concepts predict activation of a given region. Areas of SITB-related hyperactivation (precuneus/PCC and TPJ) were significantly associated with internally-oriented processes of mentalizing, semantics, and memory, while SITB-related hypoactivation (RLPFC) was significantly associated with pain, and executive control processes of task-switching, memory, and working memory (Figure S2).

#### Meta-Analyses for Types of SITBs

To test the possibility that unique abnormalities might be associated with specific types of SITBs, finer-grained meta-analyses were conducted provided sufficient power.

##### Structural Imaging Studies

Due to insufficient power, analyses could not be conducted for structural abnormalities associated with specific types of SITBs with either the stringent 20 experiment criterion or the relaxed 10 experiment criterion (Figure 1).

##### Functional Imaging Studies

No types of SITBs met the stringent 20 experiment criterion. Suicide attempts and self-harm regardless of intent (i.e., deliberate self-harm), however, met the relaxed 10 experiment criterion (Figure 2). ALE analyses yielded no significant results. MKDA with a 15mm kernel produced no significant results for suicide attempts, but revealed a significant hyperactivation for deliberate self-harm in the precuneus/PCC (x = 6, y = − 60, z = 36; Table S2), a finding consistent with the pooled analysis.

#### Moderator Analyses

To investigate whether and how differences in samples and study designs might have affected findings, we attempted to conduct moderator analyses on type of control groups, psychiatric diagnoses, study paradigms, and medication status. However, moderator analyses could not be conducted for psychiatric diagnoses due to the heterogeneous inclusion criteria among studies. Other moderator analyses were conducted when they met either the more stringent minimum of 20 experiments or the more relaxed minimum of 10 experiments.

##### Structural Imaging Studies

No structural analyses met the stringent 20 experiment criterion. There were 11 experiments from MRI studies that reported less volume in the self-injurious participants compared to psychiatric controls (Figure 1). Consistent with the overall pooled analysis, neither ALE nor MKDA yielded any significant results.

##### Functional Imaging Studies

Regarding types of control groups, separate meta-analyses were conducted for both studies that used psychiatric controls and those that used healthy controls with the minimum 10 experiment criterion. Consistent with the pooled analyses, ALE yielded no significant results. Inconsistent with the pooled analyses, MKDA with a 10mm kernel for experiments with psychiatric controls yielded significant hyperactivation in the left temporal-parietal junction (x = − 50, y = − 66, z = 28; Table S2), suggesting a moderator effect. However, this result was not replicated for MKDA using a 15mm kernel. No significant results were yielded for hypoactivation regardless of control group type.

In terms of study paradigms, a separate meta-analysis was conducted for experiments that used cognitive tasks and showed hypoactivation among self-injurious individuals. Consistent with the pooled analysis, ALE did not yield any significant findings. Inconsistent with the overall analysis, MKDA with a 15mm kernel showed significant hypoactivation in the ventral striatum (x = 2, y = 4, z = −6; Table S2), indicating a moderator effect. MKDA with a 10mm kernel did not yield significant hypoactivation. Another separate meta-analysis was conducted for experiments that used affective tasks and showed hyperactivation among self-injurious participants. ALE again did not yield any significant findings. Consistent with the pooled analysis, MKDA with a 15mm kernel produced significant hyperactivation in precuneus/PCC (x = 6, y = −54, z = 34; Table S2). This finding was not replicated for MKDA with a 10mm kernel.

With respect to medication status, no analyses met the stringent 20 experiment minimum (Figure 2). Using the 10 experiment criterion, a meta-analysis including only experiments that recruited non-medicated participants was conducted for studies that indicated hyperactivation in self-injurious individuals. Neither ALE nor MKDA yielded any significant results when only non-medicated individuals were included. A separate meta-analysis for non-medicated participants could not be conducted for studies that found hypoactivation in self-injurious individuals due to an insufficient number of studies.

## Discussion

The present study yielded four major findings: (1) existing neuroimaging research has not found consistent structural brain abnormalities between populations with and without SITBs; (2) the ALE method produced no significant findings regarding functional abnormalities, while the MKDA method produced a convergence of findings at three locations (i.e., precuneous/PCC, TPJ, and RLPFC) that have not been the focus of the SITB literature; (3) finer-grained meta-analyses for specific types of SITBs showed that deliberate self-harm might be associated with functional differences in the precuneous/PCC; and (4) moderator analyses showed checkered consistency. These findings together suggest that the extant literature provides some, but far from unanimous, support for neural correlates of SITBs. The major findings and the strengths and limitations of the present study are discussed in more detail below.

Despite previous research suggesting structural abnormalities associated with SITBs (Aguilar et al., 2008; Benedetti et al., 2011), the present meta-analysis did not yield any significant findings with either ALE or MKDA. To our surprise, the meta-analysis did not replicate previous reviews suggesting distinct structural changes associated with SITBs (e.g., van Heeringen et al., 2011; Martin et al., 2015). A primary difference between this study and previous reviews is that we subscribed to power guidelines in the field (Eickhoff et al. 2016) that required a more stringent minimum of 20 experiments for each analysis. Even though we conducted additional analyses with a more relaxed 10 experiment criterion, more studies are needed to reveal the consistent abnormalities between individuals with and without SITBs. Due to the insufficient number of studies in the literature, it is unclear whether certain structural abnormalities are associated with risk for SITBs in general or for specific types of SITBs. Additionally, we cannot rule out the possibility that individuals with SITBs might not exhibit structural abnormalities.

Our second major finding is that functional differences in the precuneous/PCC, TPJ, and RLPFC were associated with SITBs based on the MKDA method, but not the ALE method. These regions have not been a focus of the literature on SITBs. If the locations indeed reflect underlying brain abnormalities associated with SITBs, future research might benefit from examining these areas in more detail. The precuneus/PCC is one of the major nodes of default network, which is engaged during mind-wandering, and projects heavily to the memory system (Fransson & Marrelec, 2008; Greicius, Krasnow, Reiss, & Menon, 2003; Mason et al., 2007). Functional profiling of the precuneus/PCC and TPJ revealed strong associations with mentalizing including theory of mind, suggesting that individuals with SITBs may over-represent the thoughts of others. The lateral PFC is central to cognitive control (Miller & Cohen, 2001) and functional profiling of the RLPFC cluster accordingly revealed strong associations to task-switching and working memory. In the lateral PFC, progressively rostral areas are responsible increasingly temporally abstract control (Fuster, 2001; Koechlin, Ody, & Kouneiher, 2003; Nee & D’Esposito, 2016). For example, inhibition of the RLPFC results in diminished ability to plan for the future, while leaving other control processes intact (Nee & D’Esposito, 2017). Hypoactivity in the RLPFC in self-injurious individuals might suggest an ineffectiveness in controlling their behaviors with long-term goals in mind. Collectively, over-representation of the thoughts of others coupled with ineffective future-oriented control may lay a foundation for SITBs.

Even though the present meta-analysis suggests that precuneous/PCC, TPJ, and RLPFC might be involved in SITBs, future studies are needed to directly test these hypotheses. Results from the MDKA method provided evidence for functional abnormalities associated with SITBs in general, whereas the ALE method did not yield any significant findings. Given the conceptual similarities between ALE and MKDA, the lack of convergence between them in the present study leads to questions about the robustness of the findings. It is unclear if the findings from one analytic method should overweigh the lack of findings from the other method, or vice versa. Therefore, at present, it is difficult to confidently draw any conclusion about the functional abnormalities among individuals with SITBs. More studies are needed to shed light on this topic.

Our third major finding is that the MKDA method, but not the ALE method, found associations between deliberate self-harm and the precuneous/PCC. We originally intended to conduct separate meta-analyses for each type of SITBs. To our surprise, the majority of the literature focused on studying suicide attempts, with much less focus on suicide ideation, plan, and NSSI. Therefore, we were only able to perform analyses for suicide attempts and deliberate self-harm. Consistent with the pooled analysis of all SITBs, the MKDA with a 15mm kernel revealed significant hyperactivation in the precuneous/PCC for deliberate self-harm but not for suicide attempt. It is possible that brain abnormalities might be associated with general risk for SITBs instead of specific types of SITBs. Therefore, abnormalities associated with self-harm regardless of intent might be more consistent with the overall finding than only self-harm with the intent to die (i.e., suicide attempt). It is also possible that the inclusion of self-harm without intent to die simply boosted the power to detect a significant difference. Given the paucity of research on certain types of SITBs, however, it is unclear whether unique brain changes exist for specific types of SITBs or whether they are associated with a general risk for SITBs. More studies examining self-injurious phenomena other than suicide attempts are needed to provide further insight on this issue.

Lastly, moderator analyses demonstrated checkered consistency for the significant functional abnormalities yielded by the pooled meta-analyses. Even though it was within our initial intention to systematically conduct moderator analyses, the unexpectedly limited number of experiments within each moderator category prevented us from fully performing these analyses. However, within the constraints of the literature, we conducted all moderator analyses that met the more relaxed 10 experiment criterion. Significant moderator effects of type of control groups, study paradigms, and medication status were detected from the MKDA method. However, these moderator effects were not robust as the results were generated from less stringent analyses and inconsistent across MKDA and ALE. More studies are needed detect consistent moderator effects.

Taken together, the extant literature provides some support for neural correlates of SITBs but does not yet justify definitive conclusions. This study does not imply that there are no structural or functional brain abnormalities associated with SITBs or that the brain has no relevance in SITBs. Rather, the *current* state of the literature fails to unequivocally support the conclusions drawn by previous qualitative reviews. It is possible that the neural mechanisms proposed by prior reviews do play a role in SITBs, but their effects were obscured by the dearth of research in some areas and the heterogeneity among studies. Without further studies, however, our ability to draw confident conclusions on the roles of brain abnormalities in SITBs is circumscribed.

The present findings should be considered within the context of the study’s limitations. It is important to note that a meta-analysis summarizes and reflects the current state of the literature and is therefore largely constrained by the limitations of the literature. First, the statistical power of the present meta-analysis was confined both in terms of the number of experiments and participants. It was surprising that no structural findings met the new power guidelines in the field (Eickhoff et al., 2016), and few functional findings did. The small number of studies and experiments limited our ability to conduct comprehensive meta-analyses for specific types of SITBs and finer-grained moderator analyses.

Regarding sample size, It is generally well appreciated that studies with a small sample size might lack the statistical power to detect true effects; however, small sample size also reduces the likelihood that a detected result reflects a true effect (Button et al., 2013). The median sample size of the studies included in the meta-analysis is 45, which can lead to poor replicability even in one-sample tests (Turner, Paul, Miller, & Barbey, 2018), let alone two-sample tests. On the other hand, increasing sample size will only improve power if there is an underlying group-level effect to find. Recent data indicate that there are multiple neurophysiological subtypes of depression (Drysdale et al., 2017), and it is possible that SITBs are just as, or even more variable. Furthermore, group-level inferences may not apply to individuals (Fisher, Medaglia, & Jeronimus, 2018). Such data suggest that more data are needed at the individual-level. Hence, insufficient power at either the group- and/or individual-levels may contribute to the inconsistent findings in the present meta-analysis.

Second, the heterogeneity among studies in the literature might have obscured the meta-analytic findings. For example, the thresholds that studies set to control for multiple comparisons vary widely. Insufficiently corrected analyses produce false positives, adding noise to meta-analyses (Lieberman & Cunningham, 2009). Similarly, high heterogeneity exists regarding preprocessing parameters and the contrasts analyzed, which can lead to vastly different results on the same underlying data (Carp, 2012). Moreover, 43.05% of the contrasts used healthy controls instead of psychiatric controls to test for abnormalities associated with SITBs. Considering that individuals with SITBs are likely to meet diagnostic criteria for psychiatric conditions, psychiatric controls would provide a more stringent comparison and reduce the likelihood of detecting abnormalities associated with general psychopathology instead of SITBs. Further, even though the present study was unable to directly examine the effects of psychological tasks (e.g., Iowa Gambling Task, Stroop task) employed in the studies due to insufficient statistical power, it is possible that these differences contributed to the checkered consistency of the findings. Of note, heterogeneity and flexibility in study paradigms and analytical decisions might have also obscured previous reviews and meta-analyses, contributing to the mixed conclusions in the literature.

In addition to limitations of the literature, it is important to keep in mind limitations of coordinate-based meta-analysis (CBMA). CBMA has been used to identify convergent activations in numerous domains (Nee et al., 2013; Nee, Wager, & Jonides, 2007; Rottschy et al., 2012; Wager et al., 2004), but it remains an imperfect method. CBMA creates simulated statistical maps based upon peak activations reported in studies. However, the size and shape of the simulated activation clusters are unrealistic, potentially leading to both false positive and negative results. It would also be prudent to weight activation clusters by their effect size (Tench et al., 2017). However, effect size information was irregularly reported in the present sample. In an ideal scenario, meta-analysis of neuroimaging data would be performed on unthresholded statistical maps, which would at once provide size, shape, and effect size estimates. Although resources such as NeuroVault are becoming increasingly popular (Gorgolewski et al., 2015), they are not yet used widely enough to perform meta-analyses in this domain. More consistent data sharing in the future would help to determine whether the present findings were due to limitations of CBMA.

Despite the limitations, this study also demonstrates several strengths. First, even though this meta-analysis was still underpowered for some sub-analyses and moderator analyses, it demonstrates one of the largest efforts to increase power to detect true underlying effects by including neuroimaging studies on any type of SITBs. Second, the meta-analysis employed two gold-standard coordinate-based meta-analytic methods (i.e., ALE and MKDA). This signaled progress over meta-analyses that largely relied on ALE alone and allowed for evaluation of the robustness of findings. Third, we subscribed to the new power guideline in the field (Eickhoff et al., 2016), but also conducted analyses using the previous criterion for completeness (Eickhoff & Bzdok, 2013). The power standard (i.e., a minimum of 20 experiments per analysis) is considered to be more stringent than previous standards. By following these guidelines, the present study is less likely to yield spurious findings.

To summarize, the present meta-analysis aimed to evaluate the *current* empirical evidence for neural correlates of SITBs and whether it justifies any definitive conclusions about brain abnormalities among individuals with SITBs. This study conducted pooled analyses across all SITBs, separate analyses for specific types of SITBs, and moderator analyses of differences among studies. The current state of the literature failed to provide support for structural abnormalities, and provided some, yet far from unequivocal, support for functional abnormalities. The identified abnormalities in the precuneous/PCC, TPJ, and RLPFC have not been the focus of previous studies, but may offer promising future avenues of exploration. Due to the constraints of the existing literature, it is unclear whether brain abnormalities increase general risk for SITBs or unique abnormalities are associated with specific types of SITBs. Insufficient power, heterogeneity in study paradigm, flexibility in analytical decisions, and limitations of CBMA might have hindered the current study from identifying consistent and robust patterns associated with SITBs. Given the extant literature, more studies are needed to reach definitive conclusions on abnormal brain structure and function among people with a history of SITBs. Future studies should consider gathering more group and/or individual-level data, selecting stringent control groups, providing replications of previous research, and adopting standard thresholds and preprocessing parameters.

## Authorship

D.E.N and J.C.F developed the study concept. All authors contributed to the study design. X.H. and D.M.B. conducted the literature search and extracted data from published studies. K.R.M. and D.E.N. performed the data analysis. X.H. and K.R.M. interpreted the results and drafted the paper under the supervision of D.E.N. and J.C.F. All authors approved the final version of the paper for submission.

## Acknowledgements

None.

## Declaration of Interest

None.

