## Supplemental Information for "Brain Abnormalities Associated with Self-Injurious Thoughts and Behaviors: A Meta-Analysis of Neuroimaging Studies"

**Supplement 1.** List of Included Studies.

**Figure S1.** Functional Associations of Activation Sites.

**Table S1.** Description of Included Studies.

**Table S2.** Significant Results from MKDA with 10 or More Experiments.

**Supplement 1. List of Included Studies.**

Aguilar EJ, Garica-Marti G, Marti-Bonmati L, Lull JJ, Moratal D, Escarti MJ, Robles M, Gonzalez J., Guillamon MI, & Sanjuan J (2008). Left orbitofrontal and superior temporal gyrus structural changes associated to suicidal behavior in patients with schizophrenia. *Progress in Neuro-Psychopharmacology and Biological Psychiatry* 32, 1673–1676.

Amen DG, Prunella JR, Fallon JH, Amen B, & Hanks C (2009). A comparative analysis of completed suicide using high resolution brain SPECT Imaging. *The Journal of Neuropsychiatry and Clinical Neuroscience* 21, 430–439.

Audenaert K, Goethals I, Van Laere K, Lahorte P, Brans B, Versijpt J, Vervaet M, Beelaert L, Van Heeringen K, & Dierckx R (2002). SPECT neuropsychological activation procedure with the Verbal Fluency Test in attempted suicide patients. *Nuclear medicine communications* 23, 907–916.

Benedetti F, Radaelli D, Poletti S, Locatelli C, Falini A, Colombo C, & Smeraldi E (2011). Opposite effects of suicidality and lithium on gray matter volumes in bipolar depression. *Journal of Affective Disorders* 135, 139–147.

Benedetti F, Riccaboni R, Poletti S, Radaelli D, Locatelli C, Lorenzi C, Pirovano A, Smeraldi E, & Colombo C (2014). The serotonin transporter genotype modulates the relationship between early stress and adult suicidality in bipolar disorder. *Bipolar Disorders* 16, 857–866.

Besteher B, Wagner G, Koch K, Schachtzabel C, Reichenbach JR, Schlösser R, Sauer H, & Schultz CC (2016). Pronounced prefronto-temporal cortical thinning in schizophrenia: Neuroanatomical correlate of suicidal behavior? *Schizophrenia Research* 176, 151–157.

Bijttebier S, Caeyenberghs K, van den Ameele H, Achten E, Rujescu D, Titeca K, & van

Heeringen C (2015). The Vulnerability to Suicidal Behavior is Associated with Reduced Connectivity Strength. *Frontiers in Human Neuroscience* 9, 1–11.

Cao J, Chen J mei, Kuang L, Ai M, Fang W dong, Gan Y, Wang W, Chen X rong, Xu X ming, Wang H guang, & Lv Z (2015). Abnormal regional homogeneity in young adult suicide attempters with no diagnosable psychiatric disorder: A resting state functional magnetic imaging study. *Psychiatry Research - Neuroimaging* 231, 95–102.

Cao J, Chen X, Chen J, Ai M, Gan Y, Wang W, Lv Z, Zhang S, Zhang S, Wang S, Kuang L, & Fang W (2016). Resting-state functional MRI of abnormal baseline brain activity in young depressed patients with and without suicidal behavior. *Journal of Affective Disorders* 205, 252–263.

Chase HW, Segreti AM, Keller TA, Cherkassky VL, Just MA, Pan LA, & Brent DA (2017). Alterations of functional connectivity and intrinsic activity within the cingulate cortex of suicidal ideators. *Journal of Affective Disorders* 212, 78–85.

Davis TS, Mauss IB, Lumian D, Troy AS, Shallcross AJ, Zarolia P, Ford BQ, & McRae K (2014). Emotional reactivity and emotion regulation among adults with a history of self-harm: Laboratory self-report and functional MRI evidence. *Journal of Abnormal Psychology* 123, 499–509.

Ding Y, Pereira F, Hoehne A, Beaulieu M-M, Lepage M, Turecki G, & Jollant F (2016). Altered brain processing of decision-making in healthy first-degree biological relatives of suicide completers. *Molecular Psychiatry* 22, 1149–1154.

Fan T, Wu X, Yao L, & Dong J (2013). Abnormal baseline brain activity in suicidal and non-suicidal patients with major depressive disorder. *Neuroscience Letters* 534, 35–40.

Fradkin Y, Khadka S, Bessette KL, & Stevens MC (2016). The relationship of impulsivity and cortical thickness in depressed and non-depressed adolescents. *Brain Imaging and Behavior*, 1–11.

Garrion VG, Garrett A, Menon V, Weems CF, & Reiss AL (2008). Posttraumatic stress symptoms and brain function during a response-inhibition task: An fMRI study in youth. *Depression and Anxiety* 25, 514–526.

Groschwitz RC, Plener PL, Groen G, Bonenberger M, & Abler B (2016). Differential neural processing of social exclusion in adolescents with non-suicidal self-injury: An fMRI study. *Psychiatry Research - Neuroimaging* 255, 43–49.

van Heeringen K, Wu GR, Vervaet M, Vanderhasselt MA, & Baeken C (2017). Decreased resting state metabolic activity in frontopolar and parietal brain regions is associated with suicide plans in depressed individuals. *Journal of Psychiatric Research* 84, 243–248.

Hwang J-P, Lee T-W, Tsai S-J, Chen T-J, Yang C-H, Lirng J-F, & Tsai C-F (2010). Cortical and subcortical abnormalities in late-onset depression with history of suicide attempts investigated with MRI and voxel-based morphometry. *Journal of Geriatric Psychiatry and Neurology* 23, 171–184.

Jia Z, Huang X, Wu Q, Zhang T, Lui S, Zhang J, Amatya N, Kuang W, Chan RCK, Kemp GJ, Mechelli A, & Gong Q (2010). High-field magnetic resonance imaging of suicidality in patients with major depressive disorder. *American Journal of Psychiatry* 167, 1381–1390.

Johnston JAY, Wang F, Liu J, Blond BN, Wallace A, Liu J, Spencer L, Lippard ETC, Purves KL, Landeros-Weisenberger A, Hermes E, Pittman B, Zhang S, King R, Martin A, Oquendo MA, & Blumberg HP (2017). Multimodal neuroimaging of frontolimbic structure and function associated with suicide attempts in adolescents and young adults with bipolar disorder. *American Journal of Psychiatry* 174, 667–675.

Jollant F, Lawrence NS, Giampietro V, Brammer MJ, Fullana MA, Drapier D, Courtet P, & Phillips ML (2008). Orbitofrontal cortex response to angry faces in men with histories of suicide attempts. *American Journal of Psychiatry* 165, 740–748.

Jollant F, Lawrence NS, Olie E, O'Daly O, Malafosse A, Courtet P, & Phillips ML (2010). Decreased activation of lateral orbitofrontal cortex during risky choices under uncertainty is associated with disadvantageous decision-making and suicidal behavior. *NeuroImage* 51, 1275–1281.

Kang S-G, Na K-S, Choi J-W, Kim J-H, Son Y-D, & Lee YJ (2017). Resting-state functional connectivity of the amygdala in suicide attempters with major depressive disorder. *Progress in Neuro-Psychopharmacology and Biological Psychiatry* 77, 222–227.

Kim YJ, Park HJ, Jahng GH, Lee SM, Kang WS, Kim SK, Kim T, Cho AR, & Park JK (2017). A pilot study of differential brain activation to suicidal means and DNA methylation of CACNA1C gene in suicidal attempt patients. *Psychiatry Research* 255, 42–48.

Kraus A, Valerius G, Seifritz E, Ruf M, Bremner JD, Bohus M, & Schmahl C (2010). Script-driven imagery of self-injurious behavior in patients with borderline personality disorder: A pilot FMRI study. *Acta Psychiatrica Scandinavica* 121, 41–51.

Lee KH, Pluck G, Lekka N, Horton A, Wilkinson ID, & Woodruff PWR (2015). Self-harm in schizophrenia is associated with dorsolateral prefrontal and posterior cingulate activity. *Progress in Neuro-Psychopharmacology and Biological Psychiatry* 61, 18–23.

Lee SJ, Kim B, Oh D, Kim MK, Kim KH, Bang SY, Choi TK, & Lee SH (2016a). White matter alterations associated with suicide in patients with schizophrenia or schizophreniform disorder. *Psychiatry Research - Neuroimaging* 248, 23–29.

Lee YJ, Kim S, Gwak AR, Kim SJ, Kang SG, Na KS, Son YD, & Park J (2016b). Decreased regional gray matter volume in suicide attempters compared to suicide non-attempters with major depressive disorders. *Comprehensive Psychiatry* 67, 59–65.

Leyton M, Paquette V, Gravel P, Rosa-Neto P, Weston F, Diksic M, & Benkelfat C (2006). α-[11C]methyl-L-tryptophan trapping in the orbital and ventral medial prefrontal cortex of suicide attempters. *European Neuropsychopharmacology* 16, 220–223.

Mahon K, Burdick KE, Wu J, Ardekani BA, & Szeszko PR (2012). Relationship between suicidality and impulsivity in bipolar I disorder: A diffusion tensor imaging study. *Bipolar Disorders* 14, 80–89.

Marchand WR, Lee JN, Johnson S, Thatcher J, Gale P, Wood N, & Jeong EK (2012). Striatal and cortical midline circuits in major depression: Implications for suicide and symptom expression. *Progress in Neuro-Psychopharmacology and Biological Psychiatry* 36, 290–299.

Matthews S, Spadoni A, Knox K, Strigo I, & Simmons A (2012). Combat-exposed war veterans at risk for suicide show hyperactivation of prefrontal cortex and anterior cingulate during error processing. *Psychosomatic Medicine* 74, 471–475.

Minzenberg MJ, Lesh TA, Niendam TA, Cheng Y, & Carter CS (2016). Conflict-Related Anterior Cingulate Functional Connectivity Is Associated With Past Suicidal Ideation and Behavior in Recent-Onset Psychotic Major Mood Disorders. *The Journal of Neuropsychiatry and Clinical Neurosciences* 28, 299–305.

Minzenberg MJ, Lesh TA, Niendam TA, Yoon JH, Cheng Y, Rhoades RN, & Carter CS (2015). Control-related frontal-striatal function is associated with past suicidal ideation and behavior in patients with recent-onset psychotic major mood disorders. *Journal of*

*Affective Disorders* 188, 202–209.

Niedtfeld I, Kirsch P, Schulze L, Herpertz SC, Bohus M, & Schmahl C (2012). Functional connectivity of pain-mediated affect regulation in borderline personality Disorder. *PLoS ONE* 7, 1–10.

Niedtfeld I, Schulze L, Kirsch P, Herpertz SC, Bohus M, & Schmahl C (2010). Affect regulation and pain in borderline personality disorder: A possible link to the understanding of self-injury. *Biological Psychiatry* 68, 383–391.

Olié E, Jollant F, Deverdun J, de Champfleur NM, Cyprien F, Le Bars E, Mura T, Bonafé A, & Courtet P (2017). The experience of social exclusion in women with a history of suicidal acts: a neuroimaging study. *Scientific Reports* 7, 89.

Olvet DM, Peruzzo D, Thapa-Chhetry B, Sublette ME, Sullivan GM, Oquendo MA, Mann JJ, & Parsey RV (2014). A diffusion tensor imaging study of suicide attempters. *Journal of Psychiatric Research* 51, 60–67.

Oquendo MA, Placidi GPA, Malone KM, Campbell C, Keilp J, Brodsky B, Kegeles LS, Cooper TB, Parsey R V, van Heertum RL, & Mann JJ (2003). Positron emission tomography of regional brain metabolic responses to a serotonergic challenge and lethality of suicide attempts in major depression. *Archives of general psychiatry* 60, 14–22.

Osuch E, Ford K, Wrath A, Bartha R, & Neufeld R (2014). Functional MRI of pain application in youth who engaged in repetitive non-suicidal self-injury vs. psychiatric controls. . *Psychiatry Research - Neuroimaging* 223, 104–112.

Pan L, Segreti A, Almeida J, Jollant F, Lawrence N, Brent D, & Phillips M (2013a). Preserved hippocampal function during learning in the context of risk in adolescent suicide attempt. *Psychiatry Research - Neuroimaging* 211, 112–118.

Pan LA, Batezati-Alves SC, Almeida JRC, Segreti A, Akkal D, Hassel S, Lakdawala S, Brent DA, & Phillips ML (2011). Dissociable patterns of neural activity during response inhibition in depressed adolescents with and without suicidal behavior. *Journal of the American Academy of Child and Adolescent Psychiatry* 50, 602–611.

Pan LA, Hassel S, Segreti AM, Nau SA, Brent DA, & Phillips ML (2013b). Differential patterns of activity and functional connectivity in emotion processing neural circuitry to angry and happy faces in adolescents with and without suicide attempt. *Psychological Medicine* 43, 2129–2142.

Pan LA, Ramos L, Segreti AM, Brent DA, & Phillips ML (2015). Right superior temporal gyrus volume in adolescents with a history of suicide attempt. *British Journal of Psychiatry* 206, 339–340.

Peng H, Wu K, Li J, Qi H, Guo S, Chi M, Wu X, Guo Y, Yang Y, & Ning Y (2014). Increased suicide attempts in young depressed patients with abnormal temporal-parietal-limbic gray matter volume. *Journal of Affective Disorders* 165, 69–73.

Plener PL, Bubalo N, Fladung AK, Ludolph AG, & Lulé D (2012). Prone to excitement: Adolescent females with non-suicidal self-injury (NSSI) show altered cortical pattern to emotional and NSS-related material. *Psychiatry Research - Neuroimaging* 203, 146–152.

Quevedo K, Martin J, Scott H, Smyda G, & Pfeifer JH (2016). The neurobiology of self-knowledge in depressed and self-injurious youth. *Psychiatry Research - Neuroimaging* 254, 145–155.

Reitz S, Kluetsch R, Niedtfeld I, Knorz T, Lis S, Paret C, Kirsch P, Meyer-Lindenberg A, Treede RD, Baumgärtner U, Bohus M, & Schmahl C (2015). Incision and stress regulation in borderline personality disorder: Neurobiological mechanisms of self-injurious behaviour.

*British Journal of Psychiatry* 207, 165–172.

Richard-Devantoy S, Ding Y, Lepage M, Turecki G, & Jollant F (2016). Cognitive inhibition in depression and suicidal behavior: a neuroimaging study. *Psychological Medicine* 46, 933–944.

Rüsch N, Spoletini I, Wilke M, Martinotti G, Bria P, Trequattrini A, Bonaviri G, Caltagirone C, & Spalletta G (2008). Inferior frontal white matter volume and suicidality in schizophrenia. *Psychiatry Research - Neuroimaging* 164, 206–214.

Schmahl C, Bohus M, Esposito F, Treed R-D, Di Salle F, Greffrath W, Ludaescher P, Jochims A, Lieb K, Scheffler K, Hennig J, & Seifritz E (2006). Neural Correlates of Antinociception in Borderline Personality Disorder. *Archives of General Psychiatry* 63, 659–667.

Schreiner MW, Klimes-Dougan B, Mueller BA, Eberly LE, Reigstad KM, Carstedt PA, Thomas KM, Hunt RH, Lim KO, & Cullen KR (2017). Multi-modal neuroimaging of adolescents with non-suicidal self-injury: Amygdala functional connectivity. *Journal of Affective Disorders* 221, 47–55.

Taylor WD, Boyd B, McQuoid DR, Kudra K, Saleh A, & MacFall JR (2015). Widespread white matter but focal gray matter alterations in depressed individuals with thoughts of death. *Progress in Neuro-Psychopharmacology and Biological Psychiatry* 62, 22–28.

Vanyukov PM, Szanto K, Hallquist MN, Siegle GJ, Reynolds CF, Forman SD, Aizenstein HJ, & Dombrovski AY (2016). Paralimbic and lateral prefrontal encoding of reward value during intertemporal choice in attempted suicide. *Psychological Medicine* 46, 381–391.

Wagner G, Koch K, Schachtzabel C, Schultz CC, Sauer H, & Schlösser RG (2011). Structural brain alterations in patients with major depressive disorder and high risk for suicide:

Evidence for a distinct neurobiological entity? *NeuroImage* 54, 1607–1614.

Wagner G, Schultz CC, Koch K, Schachtzabel C, Sauer H, & Schlösser RG (2012). Prefrontal cortical thickness in depressed patients with high-risk for suicidal behavior. *Journal of Psychiatric Research* 46, 1449–1455.

Wallace AR (2015). Neurocircuitry of suicidal behavior in adolescents and young adults with bipolar and major depressive disorder. *Yale Medicine Thesis Digital Library*

Willeumier K, Taylor D V, & Amen DG (2011). Decreased cerebral blood flow in the limbic and prefrontal cortex using SPECT imaging in a cohort of completed suicides. *Translational Psychiatry* 1, e28.

Zhang H, Wei X, Tao H, Mwansisya TE, Pu W, He Z, Hu A, Xu L, Liu Z, Shan B, & Xue Z (2013). Opposite effective connectivity in the posterior cingulate and medial prefrontal cortex between first-episode schizophrenic patients with suicide risk and healthy controls. *PLoS ONE* 8, 1–8.

Zhang S, Chen J, Kuang L, Cao J, Zhang H, Ai M, Wang W, Zhang S, Wang S, Liu S, & Fang W (2016). Association between abnormal default mode network activity and suicidality in depressed adolescents. *BMC Psychiatry* 16, 337.

**Figure S1. Functional Associations of Activation Sites.**

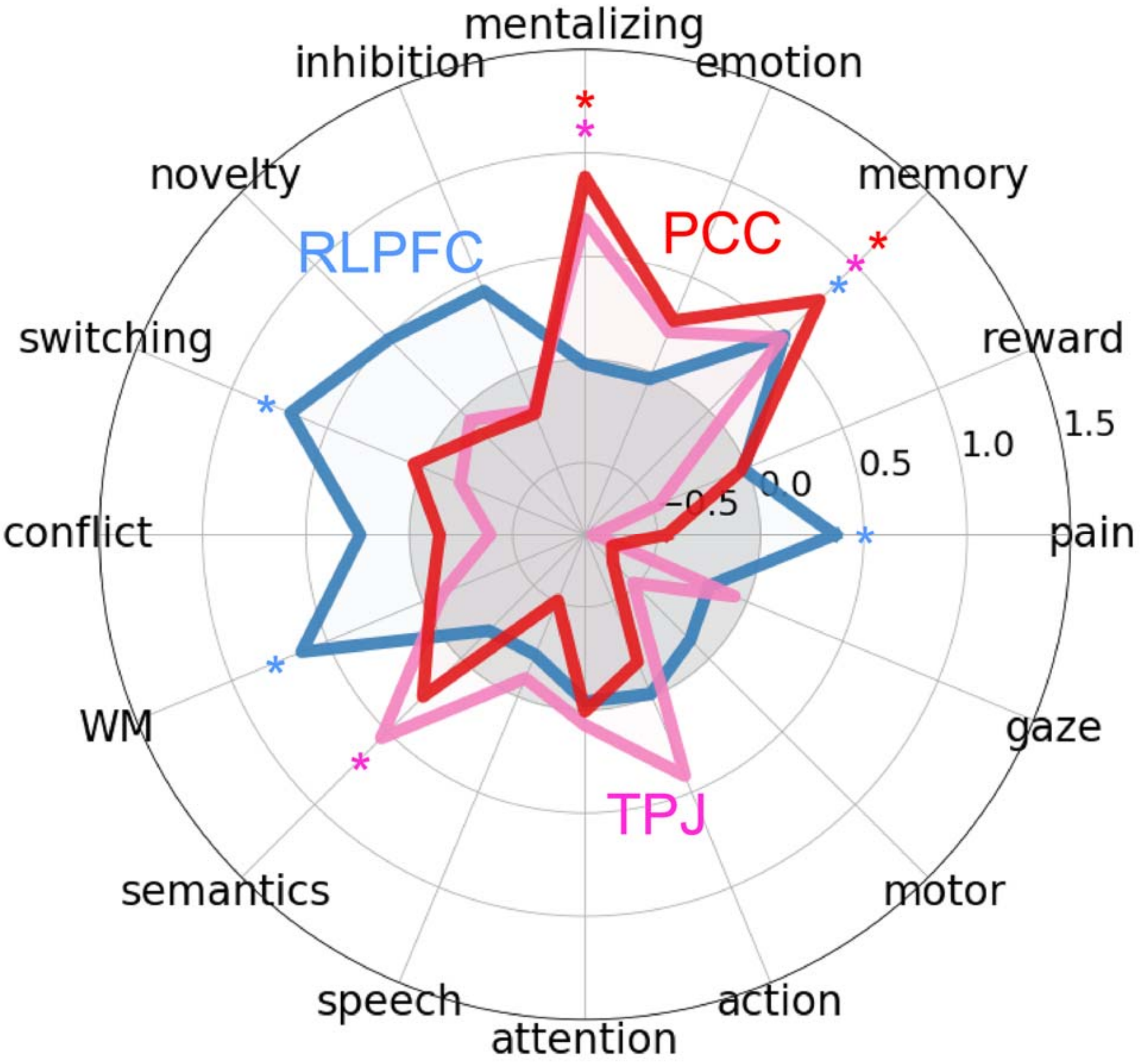


*Note*. Each cluster is illustrated to show which psychological concepts best predicted its activation based upon methods of de la Vega et al. (2017). Strength of association is measured in the log odds-ratio. Permutation-based significance after correction for multiple comparisons via false-discovery rate at q < 0.01 is indicated by an asterisk.

**Table S1. Description of Included Studies.**

| Paper | Imaging Techniques | SITB Group N | Control Group N | Control Type | Medication | Sample Age | Tasks | SITBs Type | Number of Contrasts | Source of Coordinates |
|---|---|---|---|---|---|---|---|---|---|---|
| Aguilar et al., 2008 | MRI | 13 | 24 | Psychiatric | Unclear | Adults Only | NA | Suicide Attempt | 2 | Table 2 |
| Amen et al., 2009 | SPECT | 9 | 12 | Healthy | Some Medicated | Adults Only | Connor's Continuous Performance Test | Suicide Death | 15 | Table 2 |
| | | 12 | 12 | Healthy | | | | | 25 | Table 3 |
| | | 9 | 12 | Psychiatric | | | | | 6 | Table 6 |
| | | 12 | 12 | Psychiatric | | | | | 15 | Table 7 |
| Audenaert et al., 2002 | SPECT | 20 | 20 | Healthy | Some Medicated | Adults Only | Controlled Oral Word Association Test | Suicide Attempt | 16 | Figures 1-3 |
| Benedetti et al., 2011 | MRI | 19 | 38 | Psychiatric | Some Medicated | Adults Only | NA | Suicide Attempt | 40 | Table 2 |
| Benedetti et al., 2014 | MRI | 32 | 104 | Psychiatric | Some Medicated | Adults Only | NA | Suicide Attempt | 1 | Figure 3 |
| Besteher et al., 2016 | MRI | 14 | 23 | Psychiatric | All Medicated | Adults Only | NA | Suicide Attempt | 1 | Table 3 |
| Bijttebier et al., 2015 | DTI | 13 | 15 | Psychiatric | Some Medicated | Adults Only | NA | Suicide Attempt | 4 | Table 2 |
| | | 13 | 22 | Healthy | | | | | 4 | |
| Cao et al., 2015 | fMRI | 19 | 20 | Healthy | Unclear | Adults Only | NA | Suicide Attempt | 14 | Table 2 |
| Cao et al., 2016 [a] | fMRI | 35 | 18 | Psychiatric | Some Medicated | Adolescents and Adults | NA | Suicide Attempt | 6 | Table 2 |
| | | 35 | 47 | Healthy | | | | | 4 | |
| Chase et al., 2017 | fMRI | 34 | 40 | Healthy | All Medicated | Adults Only | NA | Suicide Ideation | 2 | Figure 2 |
| | | 18 | 16 | Self-Injurious | | | | Suicide Attempt | 2 | |
| Davis et al., 2014 | fMRI | 21 | 27 | Healthy | Unclear | Adults Only | Negative Emotional Reactivity and Regulation Task | Self-Harm (Regardless of Intent) | 6 | Table 5 |
| Ding et al., 2016 | fMRI | 17 | 16 | Psychiatric | None Medicated | Adults only | Iowa Gambling Task | Suicide Death | 9 | Table 2 |
| | | 17 | 19 | Healthy | | | | | | |
| Fan et al., 2013 | fMRI | 27 | 9 | Psychiatric | Some Medicated | Adults Only | NA | Suicide Attempt | 2 | Table 2 |
| | | 27 | 57 | Healthy | | | | | 5 | |
| Fradkin et al., 2016 | MRI | 29 | 29 | Healthy | Some Medicated | Adolescents and Adults | NA | Suicide Attempt | 4 | Table 2 |
| Garrion et al., 2008 | fMRI | 7 | 9 | Psychiatric | None Medicated | Adolescents Only | Go/No-Go Task | Self-Harm (Regardless of Intent) | 4 | Table 5 |

**Table S1 - continued**

| Paper | Imaging Techniques | SITB Group N | Control Group N | Control Type | Medication | Sample Age | Tasks | SITBs Type | Number of Contrasts | Source of Coordinates |
|---|---|---|---|---|---|---|---|---|---|---|
| Groschwitz et al., 2016 | fMRI | 14 | 14 | Psychiatric | Some Medicated | Adolescents Only | Cyberball Game | NSSI | 3 | Table 2 |
| | | | | | | | | | 3 | Table 3 |
| Hwang et al., 2010 | MRI | 27 | 43 | Psychiatric | Unclear | Elderly Only | NA | Suicide Attempt | 54 | Tables 3 & 4 |
| Jia et al., 2010 | DTI | 16 | 36 | Psychiatric | None Medicated | Adults Only | NA | Suicide Attempt | 2 | Table 2 |
| | | 16 | 52 | Healthy | | | | | 2 | |
| Johnston et al., 2017 | fMRI | 26 | 42 | Psychiatric | Some Medicated | Adolescents and Adults | Emotion Face Processing Task | Suicide Attempt | 4 | Table 2 |
| | MRI | | | | | | NA | | 3 | |
| | DTI | | | | | | NA | | 3 | |
| Jollant et al., 2008 | fMRI | 13 | 14 | Psychiatric | Unclear | Adults Only | Emotion Face Viewing Task | Suicide Attempt | 13 | Table 3 |
| Jollant et al., 2010 | fMRI | 13 | 12 | Psychiatric | Some Medicated | Adults Only | Iowa Gambling Task | Suicide Attempt | 3 | Page 1278 |
| Kang et al., 2017 [b] | fMRI | 19 | 19 | Psychiatric | Some Medicated | Adults Only | NA | Suicide Attempt | 3 | Table 1 |
| Kim et al., 2017 | fMRI | 14 | 22 | Psychiatric | Unclear | Adults Only | Viewing pictures of faces, suicidal means, and natural landscape | Suicide Attempt | 4 | Figure 2 |
| Kraus et al., 2010 | fMRI | 11 | 10 | Healthy | None Medicated | Adults Only | Script-Driven Imagery of Self-Injurious Behaviors | Self-Harm (Regardless of Intent) | 2 | Page 46 |
| Lee et al., 2015 | fMRI | 14 | 14 | Psychiatric | Unclear | Adults Only | Go/No-Go Task | Self-Harm (Regardless of Intent) | 6 | Table 3 |
| Lee, Kim, Gwak et al., 2016 [b] | MRI | 19 | 19 | Psychiatric | Some Medicated | Adults Only | NA | Suicide Attempt | 2 | Table 2 |
| Lee, Kim, Oh et al., 2016 | DTI | 15 | 41 | Psychiatric | All Medicated | Adults Only | NA | Suicide Attempt | 1 | Page 25 |
| Leyton et al., 2006 | PET | 10 | 16 | Healthy | None Medicated | Adults Only | NA | Suicide Attempt | 5 | Page 222 |
| Mahon et al., 2012 | DTI | 14 | 15 | Psychiatric | Some Medicated | Adults Only | NA | Suicide Attempt | 2 | Figure 1 |

**Table S1 - continued**

| Paper | Imaging Techniques | SITB Group N | Control Group N | Control Type | Medication | Sample Age | Tasks | SITBs Type | Number of Contrasts | Source of Coordinates |
|---|---|---|---|---|---|---|---|---|---|---|
| Marchand et al., 2012 | fMRI | 6 | 16 | Psychiatric | None Medicated | Adults Only | Self-Paced Button Pressing Task | Self-Harm (Regardless of Intent) | 14 | Tables 2, 3, 5 |
| | | 13 | 9 | Psychiatric | | | | Suicide Ideation | 7 | Tables 2 & 4 |
| Matthews et al., 2012 | fMRI | 13 | 13 | Psychiatric | Some Medicated | Adults Only | Stop Task | Suicide Ideation | 12 | Table 3 |
| Minzenberg et al., 2015 [c] | fMRI | 16 | 14 | Psychiatric | All Medicated | Adults Only | Continuous Performance Task (AX Version) | Suicide Ideation | 3 | Table 2 |
| | | 8 | 8 | Self-Injurious | | | | All Suicidal Behaviors | 6 | |
| Minzenberg et al., 2016 [c] | fMRI | 16 | 14 | Psychiatric | All Medicated | Adults Only | Continuous Performance Task (AX Version) | Suicide Ideation | 24 | Table 2 |
| | | | | | | | | Suicide Ideation & Plan | 10 | |
| | | | | | | | | All Suicidal Behaviors | 11 | |
| Niedtfeld et al., 2010 [d] | fMRI | 20 | 23 | Healthy | None Medicated | Adults Only | Viewing Negative and Neutral Valence Pictures and Receiving Thermal Stimuli | Self-Harm (Regardless of Intent) | 51 | Tables 1 & 3 |
| Niedtfeld et al., 2012 [d] | fMRI | 20 | 23 | Healthy | None Medicated | Adults Only | Viewing Negative and Neutral Valence Pictures and Receiving Thermal Stimuli | Self-Harm (Regardless of Intent) | 23 | Tables S1-2, & S4-6 |
| | | | | | | | | | 4 | Tables S2, S3, & S6 |
| | | | | | | | | | 9 | Tables S1, S3-6 |
| Olié et al., 2017 | fMRI | 36 | 41 | Psychiatric | Some Medicated | Adults Only | Cyberball Game | Suicide Attempt | 2 | Figure 2 |
| Oquendo et al., 2003 | PET | 16 | 9 | Self-Injurious | None Medicated | Adults Only | NA | Suicide Attempt | 9 | Figure 1 |
| Olvet et al., 2014 | DTI | 13 | 39 | Psychiatric | Some Medicated | Adults Only | NA | Suicide Attempt | 1 | Figure 1 |

**Table S1 - continued**

| Paper | Imaging Techniques | SITB Group N | Control Group N | Control Type | Medication | Sample Age | Tasks | SITBs Type | Number of Contrasts | Source of Coordinates |
|---|---|---|---|---|---|---|---|---|---|---|
| Osuch et al., 2014 | fMRI | 13 | 15 | Psychiatric | Some Medicated | Adolescents and Adults | Self-administered and Experimenter-administered cold stimuli | NSSI | 35 | Tables 2 & 3, Figure 5 |
| Pan et al., 2011 [e] | fMRI | 15 | 15 | Psychiatric | Some Medicated | Adolescents Only | Go/No-Go Task | Suicide Attempt | 3 | Table 2 |
| Pan et al., 2015 | MRI | 28 | 31 | Psychiatric | Some Medicated | Adolescents Only | NA | Suicide Attempt | 3 | Table DS2 |
| | | 28 | 41 | Healthy | | | | | 13 | |
| Pan, Hassel et al., 2013 [e] | fMRI | 14 | 15 | Psychiatric | Some Medicated | Adolescents Only | Emotion Face Viewing Task | Suicide Attempt | 22 | Tables 3-5 |
| | | 14 | 15 | Healthy | | | | | 17 | |
| Pan, Segreti, et al., 2013 [e] | fMRI | 15 | 14 | Psychiatric | Some Medicated | Adolescents Only | Iowa Gambling Task | Suicide Attempt | 6 | Tables 2 & 3 |
| | | 15 | 13 | Healthy | | | | | 7 | |
| Peng et al., 2014 | MRI | 20 | 18 | Psychiatric | All Medicated | Adults Only | NA | Suicide Attempt | 1 | Table 2 |
| | | 20 | 28 | Healthy | | | | | 2 | |
| Plener et al., 2012 | fMRI | 9 | 9 | Healthy | Some Medicated | Adolescents Only | Viewing Emotional IAPS Pictures and NSSI Pictures | NSSI | 15 | Table 3 |
| Quevedo et al., 2016 | fMRI | 50 | 36 | Psychiatric | Some Medicated | Adolescents Only | Interpersonal Self-Processing task | NSSI | 8 | Table 2 |
| | | 50 | 37 | Healthy | | | | | 2 | |
| Reitz et al., 2015 | fMRI | 21 | 17 | Healthy | None Medicated | Adults Only | Montreal Imaging Stress Task and Pain Induction | NSSI | 3 | Table 2 |
| | | | | | | | | | 1 | |
| | | | | | | | | | 2 | |
| Richard-Devantoy et al., 2016 | fMRI | 26 | 23 | Psychiatric | Some Medicated | Adults Only | Go/No-Go Task | Suicide Attempt | 1 | Supplement |
| | | | 28 | Healthy | | | | | 1 | |
| Rüsch et al., 2008 | MRI | 10 | 45 | Psychiatric | Some Medicated | Adults Only | NA | Suicide Attempt | 2 | Page 209 |
| Schmahl et al., 2006 | fMRI | 12 | 12 | Healthy | None Medicated | Adults Only | Thermal Stimuli | Self-Harm (Regardless of Intent) | 6 | Table 3 |
| Schreiner, 2017 | fMRI | 25 | 20 | Healthy | Some Medicated | Adolescents and Adults | NA | NSSI | 8 | Table 2 |
| | | 24 | 17 | Healthy | | | Emotion Face Matching Task | | 15 | Tables 3, S1, S2 |

**Table S1 - continued**

| Paper | Imaging Techniques | SITB Group N | Control Group N | Control Type | Medication | Sample Age | Tasks | SITBs Type | Number of Contrasts | Source of Coordinates |
|---|---|---|---|---|---|---|---|---|---|---|
| Taylor et al., 2015 | DTI | 21 | 53 | Psychiatric | Unclear | Adults Only | NA | Suicide Ideation | 2 | Table 2 |
| | DTI | 21 | 91 | Healthy | | | | | 4 | |
| | MRI | 21 | 53 | Psychiatric | | | | | 4 | Figure 1 |
| Van Heeringen et al., 2017 | PET | 17 | 20 | Healthy | All Medicated | Adults Only | NA | Suicide Ideation | 3 | Table 2 |
| | | | | | | | | Suicide Plan | 5 | |
| Vanyukov et al., 2016 | fMRI | 13 | 22 | Healthy | Unclear | Adults and Elderly | Delay Discounting Task | Suicide Attempt | 1 | Figure 2 |
| Wagner et al., 2011 [f] | MRI | 15 | 15 | Clinical | None Medicated | Adults Only | NA | Suicide Risk | 2 | Page 1610 |
| Wagner et al., 2012 [f] | MRI | 15 | 15 | Clinical | None Medicated | Adults Only | NA | Suicide Risk | 2 | Page 1452 |
| Wallace 2015 | DTI | 21 | 25 | Psychiatric | Some Medicated | Adolescents and Adults | NA | Suicide Attempt | 3 | Table 4 |
| | | 18 | 17 | Psychiatric | | | | | 1 | Table 5 |
| | | 39 | 42 | Psychiatric | | | | | 6 | Table 3 |
| | | 21 | 43 | Healthy | | | | | 3 | |
| | | 18 | 43 | Healthy | | | | | 1 | |
| | | 39 | 43 | Healthy | | | | | 6 | |
| Willeumier et al., 2011 | SPECT | 21 | 36 | Psychiatric | None Medicated | Adolescents and Adults | NA | Suicide Death | 10 | Table 3 |
| | | 21 | 27 | Healthy | | | | | 10 | Table 4 |
| Zhang et al., 2013 | fMRI | 14 | 19 | Clinical | Unclear | Adults Only | 2-Back Task | Suicide Risk | 2 | Page 3 |
| | | 14 | 15 | General | | | | | | |
| Zhang et al., 2016 [a] | fMRI | 35 | 18 | Psychiatric | Unclear | Adolescents and Adults | NA | Suicide Attempt | 3 | Table 2 |
| | | 35 | 47 | Healthy | | | | | 5 | |

*Note.* Papers with the same superscript letter used the same subject group.

**Table S2. Significant Results from MKDA with 10 or More Experiments.**

| Radius Size | Activation | Study Type | Patient Type | Control Type | Peak Coordinate (MNI) | Number of voxels | Number of foci | Number of subjects |
|---|---|---|---|---|---|---|---|---|
| **10mm** | **Hyperactivation** | Emotional Tasks | SITBs | All Controls | 6,-58,30 | 415 | 111 | 98 |
| | | All Functional Studies | SITBs | All Controls | 6,-58,34 | 233 | 227 | 305 |
| | | All Functional Studies | SITBs | PSY | -50,-66,28 | 414 | 93 | 116 |
| **15mm** | **Hypoactivation** | Cognitive Tasks | SITBs | All Controls | 2,4,-6 | 778 | 80 | 106 |
| | | All Functional Studies | SITBs | All Controls | 36,50,18 | 669 | 157 | 306 |
| | **Hyperactivation** | All Functional Studies | SITBs | All Controls | 4,-56,32<br>48,-62,24 | 1249<br>445 | 227 | 305 |
| | | All Functional Studies | SA, Self-Harm behavior, & NSSI | All Controls | 6,-60,36 | 439 | 109 | 187 |
| | | Emotional Tasks | SITBs | All Controls | 6,-54,34 | 1519 | 111 | 98 |

*Note*. PSY: Matched psychiatric populations; SITBs: all self-injurious thoughts and behaviors; SA: patients with suicide attempts; NSSI: patients with non-suicidal self-injury.
